# Towards discrimination of mammary pathogenic *Escherichia coli* (MPEC) in cattle based on possession of different iron acquisition systems

**DOI:** 10.1101/2021.08.31.458473

**Authors:** Hamideh Kalateh Rahmani, Gholamreza Hashemi Tabar, Mahdi Askari Badouei, Babak Khoramian

## Abstract

Most efforts to elucidate virulence mechanisms of mammary pathogenic *Escherichia coli* (MPEC), causative agent of bovine clinical mastitis, have been failed but some recent studies introduced iron acquisition systems as major role players in pathogenicity. Here, we investigated the different iron uptake systems genotypes and assessed how they relate to virulence potential of MPEC. In total, 217 *E. coli* isolates (MPEC= 157, fecal isolates= 60) were screened for the presence of nine genes related to iron acquisition (*iroN, iutA, fecA, fyuA, sitA, irp2, iucD, chuA* and *tonB*) and phylogenetic groups were also determined. Next, bacterial growth potential and survival in raw and UHT milk which are representative for crucial steps in mastitis development were evaluated. In addition, the mineral consumption of *E. coli* cultured in milk were measured. The results showed that MPEC strains considerably tend to possess *fecA* (93%, *p*= 0.000) and belong to phylogenetic group A (42%, *p*= 0.042). The *fecA*+ strains from both mastitis and fecal *E. coli* had a significant (*p*= 0.000) growth potential in raw and UHT milk. Interestingly, for the first time, it was shown that *fecA*+ isolates consumed less amounts of iron and other metal ions. Overall, it seems that the uptake systems related to *fecA* contributes to overcoming harsh conditions of milk and genetic lineages could also affect pathogenicity of MPEC. These findings could lead us to define MPEC with more clarity based on genotypes or growth potential in milk and possibly promote novel solutions to control mastitis more effectively in future.

**Importance:** Mastitis is one of the most costly concerns in bovine medicine and the main cause of antibiotic use in dairy herds’ worldwide. As a rule of thumb, it was believed that extraintestinal pathogenic *E. coli* expands iron acquisition virulence arsenal to enhance pathogenic potential in the environments of the host outside the intestines. The present study indicated that the long believed idea of possession of diverse mineral acquisition systems and siderophores in all ExPEC groups could be a fairy tale. Along with recent studies, the present research showed that the *fec* operon could be the minimal necessary factor to overcome the harsh conditions of milk with limiting mineral concentrations. Obviously, the *fecA*+ isolates were fast-growing and consumed less amounts of minerals. It seems that the *fec* locus and its related metabolic pathways could be the potential targets for diagnostic, preventive, and therapeutic purposes.

## Introduction

Pathogenic *Escherichia coli* cause various enteric and extraintestinal infections in animals and human. Although the main virulence elements of diarrheagenic *E. coli* (DEC) and their role in pathogenesis of enteric infections have been clearly defined, we know less about such traits in extraintestinal pathogenic *E. coli* (ExPEC), in particular, most attempts towards defining specific virulence factors of mammary pathogenic *E. coli* (MPEC) have been unsuccessful. Therefore, some researchers questioned the classification of mastitis causing *E. coli* into a separate pathotype, MPEC (1). However, few studies which mostly benefited from whole genome sequencing data introduced few candidate genetic elements in mastitis causing *E. coli* which seem to be necessary for development of mastitis. The most promising results for discrimination of MPEC pathotype has been obtained by few recent studies that showed the importance of iron acquisition systems in pathogenicity of MPEC. In these examples, the *fec* locus which is responsible for iron acquisition through ferric citrate has been incriminated to play a major role in pathogenicity (2–4). Noticeable studies have been conducted during recent years, focusing on the relationship between *fecA* (ferric citrate outer membrane receptor) and MPEC. However, such hypothesis needs to be verified through further studies on clinical isolates.

Although the incremental effect of iron in severity of infection and positive influence of this factor on elevating the growth of *E. coli* in milk have been shown long before recent studies (5, 6), confirming the exact relations among iron, MPEC and milk were not possible due to the complexity of these systems. There are various aspects of host-pathogen relationship that should be considered when assessing MPEC pathogenicity. First, milk is a poor source of iron and contains different bonded forms of this mineral (lactoferrin, transferrin, etc.) which mostly have low bioavailability (7). Moreover, there are numerous specific and non-specific iron uptake systems in *E. coli* including different siderophores, iron uptake through haem, ferric citrate uptake and ferrous ion uptake that challenge the investigation of iron uptake systems (8,9). Among mentioned systems, siderophores have been widely studied and are known as the main mechanism of ferric (Fe^3+^) acquisition in both pathogenic and non-pathogenic *E. coli*. There are four famous siderophore molecules (enterobactin, salmochelin, yersiniabactin and aerobactin) in *E. coli*. It seems that most of the *E. coli* isolates have the ability to gain iron via enterobactin, while a smaller proportion of *E. coli* isolates can produce other siderophores (10). In comparison to siderophores, less attention has been paid to other iron uptake systems.

In the lights of recent genomic findings, the goals of the current study were to investigate the effects of iron acquisition systems on mastitis causing *E. coli* and assessing the possibility of a specific genotypic and/or phenotypic characteristic as an indicator for MPEC pathotype. For these purposes, first we determined genotypes of various iron acquisition systems and compared the data with fecal isolates’ genotypes. As there are extensive iron acquisition mechanisms and genes, we focused on some of the most well-known genes related to iron uptake which have a proven role in virulence of *E. coli* (siderophores including salmochelin, yersiniabactin and aerobactin; ferric citrate uptake; iron uptake through haem; energy transducer for iron uptake systems). After that, we tried to elucidate the possible relations between genotypes and phenotypes. In other words, we compared different genotypes with particular phenotypic features including growth capacity and survival in bovine milk and mineral consumption to validate if any genotypes could reflect the pathogenic potential.

## Results

### Genetic profiles of iron acquisition systems

The current study assessed the different genetic elements related to iron acquisition as possible discrimination factors for MPEC. Thus, fecal isolates of *E. coli* from the same farms as MPEC strains isolated were chosen to be compared to MPEC genotypes. A total of 217 *E. coli* isolates (MPEC= 157, fecal= 60) were screened for presence of *iroN, iutA, fecA, fyuA, sitA, irp2, iucD, chuA*, and *tonB*. The genes *tonB* followed by *fecA* were the most prevalent genes in MPEC (99.4% and 93%, respectively), fecal isolates (98.3% and 50%, respectively), and in total (99.1% and 81.1%, respectively). In contrast, *iucD* was the least frequent gene in total (9.7%).

In the MPEC isolates, *chuA* was detected as the least frequent gene (8.3%). Moreover, genes responsible for iron acquisition through aerobactin siderophore (*iutA* and *iucD*) were only found in 6.7% of fecal isolates. By applying Fisher’s exact test, only distribution of *fecA* gene achieved statistical significance (*p*= 0.000) which means it was more likely to be found in MPEC rather than fecal isolates. Other genes showed no significant tendency to be present in MPEC or fecal isolates. Table 1 presents the results in details.

**Table 1.**
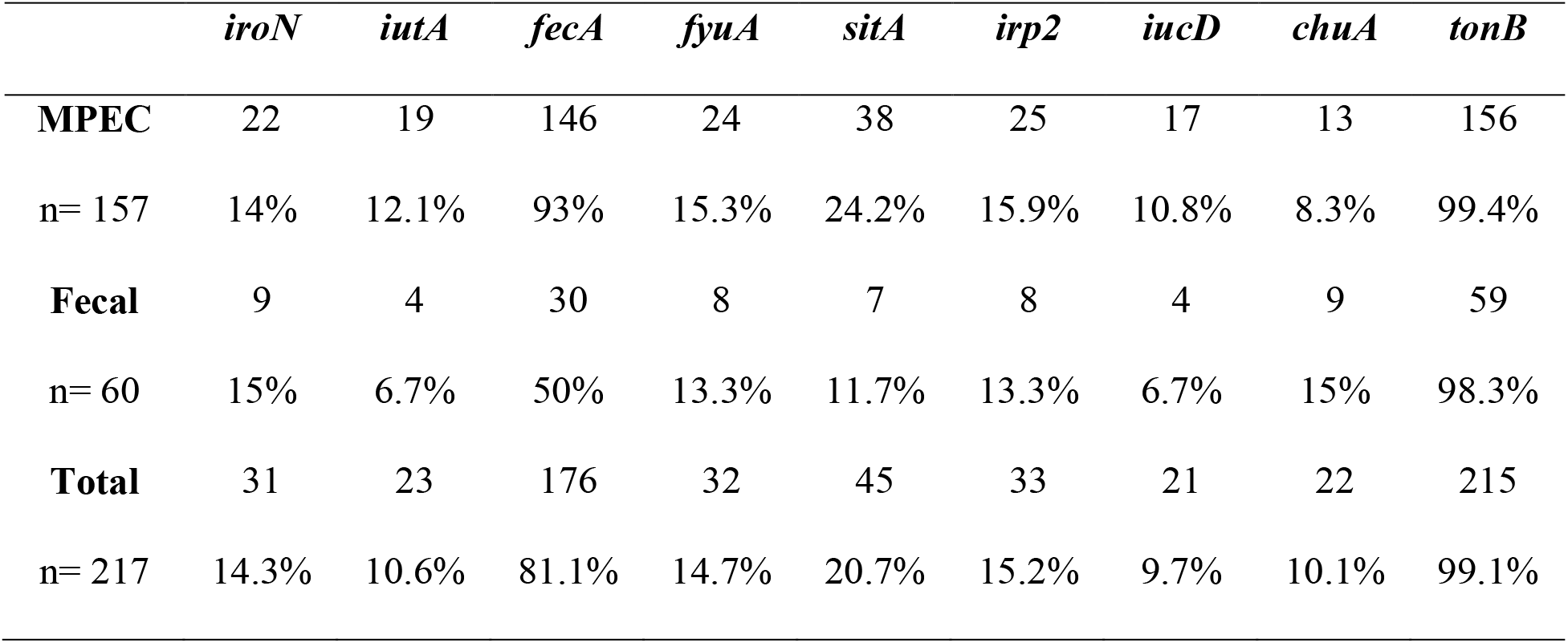
Frequency of genes related to iron acquisition in MPEC and fecal *E. coli* isolates (n, %)

Since various iron uptake systems could work together, we also compared the genetic patterns (combination of detected genes) of the isolates. Based on possessing each of the nine genes, MPEC and fecal isolates were categorized into 26 and 17 genetic profiles, respectively. Among them, only 12 patterns shared between isolates from different sources which comprised 84.2% of all isolates (82.1% of MPEC and 90.1% of fecal isolates). The patterns *fecA_tonB* (98/157 isolates, 62.4%) and *tonB* (21/60 isolates, 35%) were the most prevalent patterns in MPEC and fecal isolates, respectively. The pattern *fecA_tonB* (110/217 isolates, 50.69%) was also the most frequent pattern in total. Table 2 represents the distribution of isolates among genetic patterns in details.

**Table 2.** Distribution of isolates in genetic patterns in MPEC and fecal *E. coli* isolates (n, %)

| Genetic Patterns | MPEC<br>n=157 | Fecal<br>n= 60 | Total<br>n= 217 | p-value |
| --- | --- | --- | --- | --- |
| *fecA* | - | 1 (1.66%) | 1 (0.46%) | 0.276 |
| *tonB* | 3 (1.9%) | 21 (35%) | 24 (11.05%) | **\*0.000** |
| *iroN_fecA* | 1 (0.63%) | - | 1 (0.46%) | 1.000 |
| *iroN_tonB* | 1 (0.63%) | - | 1 (0.46%) | 1.000 |
| *fecA_tonB* | 98 (62.42%) | 12 (20%) | 110 (50.69%) | **\*0.000** |
| *fyuA_tonB* | - | 1 (1.66%) | 1 (0.46%) | 0.276 |
| *sitA_tonB* | 1 (0.63%) | 1 (1.66%) | 2 (0.92%) | 0.477 |
| *irp2_tonB* | - | 1 (1.66%) | 1 (0.46%) | 0.276 |
| *chuA_tonB* | 3 (1.9%) | 3 (5%) | 6 (2.76%) | 0.351 |
| *iroN_fecA_tonB* | 2 (1.27%) | 7 (11.66%) | 9 (4.14%) | **\*0.002** |
| *fecA_sitA_tonB* | 6 (3.82%) | - | 6 (2.76%) | 0.190 |
| *fecA_chuA_tonB* | 2 (1.27%) | 2 (3.33%) | 4 (1.84%) | 0.306 |
| *fyuA_irp2_tonB* | - | 2 (3.33%) | 2 (0.92%) | 0.076 |
| *sitA_chuA_tonB* | - | 1 (1.66%) | 1 (0.46%) | 0.276 |
| *iroN_fecA_sitA_tonB* | 2 (1.27%) | - | 2 (0.92%) | 1.000 |
| *iutA_fecA_sitA_tonB* | 1 (0.63%) | - | 1 (0.46%) | 1.000 |
| *iutA_fecA_iucD_tonB* | 1 (0.63%) | - | 1 (0.46%) | 1.000 |
| *fecA_fyuA_irp2_tonB* | 5 (3.18%) | 2 (3.33%) | 7 (3.22%) | 1.000 |
| *fecA_sitA_chuA_tonB* | 3 (1.9%) | 1 (1.66%) | 4 (1.84%) | 1.000 |
| *iroN_iutA_sitA_irp2_tonB* | 1 (0.63%) | - | 1 (0.46%) | 1.000 |
| *iroN_fecA_fyuA_irp2_tonB* | 3 (1.9%) | 1 (1.66%) | 4 (1.84%) | 1.000 |
| *iroN_fyuA_sitA_irp2_tonB* | 1 (0.63%) | - | 1 (0.46%) | 1.000 |
| *iutA_fecA_sitA_iucD_tonB* | 3 (1.9%) | 1 (1.66%) | 4 (1.84%) | 1.000 |
| *fecA_fyuA_sitA_irp2_tonB* | 2 (1.27%) | - | 2 (0.92%) | 1.000 |
| *iroN_iuA_fecA_sitA_iucD_tonB* | 5 (3.18%) | 1 (1.66%) | 6 (2.76%) | 1.000 |
| *iroN_fecA_fyuA_sitA_irp2_tonB* | 2 (1.27%) | - | 2 (0.92%) | 1.000 |
| *fecA_fyuA_sitA_irp2_chuA_tonB* | 3 (1.9%) | - | 3 (1.38%) | 0.563 |
| *iutA_fecA_fyuA_sitA_irp2_iucD_tonB* | 2 (1.27%) | - | 2 (0.92%) | 1.000 |
| *iutA_fyuA_sitA_irp2_iucD_chuA_tonB* | 1 (0.63%) | - | 1 (0.46%) | 1.000 |
| *iroN_iutA_fecA_fyuA_sitA_irp2_iucD_tonB* | 4 (2.54%) | - | 4 (1.84%) | 0.578 |
| *iutA_fecA_fyuA_sitA_irp2_iucD_chuA_tonB* | 1 (0.63%) | 2 (3.33%) | 3 (1.38%) | 0.186 |

\*significant difference

Statistical analysis revealed a significant difference (*p*= 0.000) in distribution of genetic patterns in MPEC and fecal groups. Specifically, it also showed that fecal isolates are more likely to be involved in patterns *tonB* (*p*= 0.000) and *iroN_fecA_tonB* (*p*= 0.002); on the contrary, MPEC isolates were more associated with the *fecA_tonB* pattern (*p*= 0.000).

During the screening, no isolate was found positive for all of the nine genes. Thus, isolates were categorized in to eight groups based on possession of one to eight iron acquisition genes (IAG) designated as IAG 1 to IAG 9. Generally, there was a considerable difference (*p*= 0.000) in distribution of MPEC and fecal isolates in different groups (table 3). The majority of the isolates were placed in low number groups. A total of 117 MPEC (74.5%) and 52 fecal (86.6%) isolates scored IAG1 to IAG3. Moreover, MPEC isolates preferred combination of two genes (*p*= 0.000), while fecal isolates were more prevalent in singleton (*p*= 0.000) and three-gene (*p*= 0.005) groups in comparison with MPEC.

**Table 3.** Iron acquisition genes possessions (IAG) scores in MPEC and fecal *E. coli* isolates (n, %)

|  | Number of IAG |  |  |  |  |  |  |  |
| --- | --- | --- | --- | --- | --- | --- | --- | --- |
|  | 1 | 2 | 3 | 4 | 5 | 6 | 7 | 8 |
| **MPEC** | 3 | 104 | 10 | 12 | 10 | 10 | 3 | 5 |
| n= 157 | 1.9% | 66.2% | 6.4% | 7.6% | 6.4% | 6.4% | 1.9% | 3.2% |
| **Fecal** | 22 | 18 | 12 | 3 | 2 | 1 | 0 | 2 |
| n= 60 | 36.7% | 30% | 20% | 5% | 3.3% | 1.7% | 0% | 3.3% |
| **Total** | 25 | 122 | 22 | 15 | 12 | 11 | 3 | 7 |
| n= 217 | 11.5% | 56.2% | 10.1% | 6.9% | 5.5% | 5.1% | 1.4% | 3.2% |
| **p-value** | **0.000** | **0.000** | **0.005** | 0.765 | 0.518 | 0.297 | 0.563 | 1.000 |

### Phylogenetic groups

Isolates were categorized into six phylogenetic groups A, B1, B2, C, D, E and an unknown group. The distribution of phylogroups in the two sets of isolates (MPEC and fecal) was significantly different (*p* = 0.041). Moreover, the prevalence of groups A and B1 showed significant differences (*p*= 0.042, *p*= 0.022, respectively). Infact, phylogroup A was the most common group in MPEC (42%), and B1 was more prevalent among fecal members (58.3%). Furthermore, no fecal isolate was assigned to B2 and unknown groups.

Evaluation of phylogroups and iron uptake genes revealed that members of group E were the poorest isolates in terms of possessing iron acquisition genes. All its members lacked *iroN, iutA, fyuA, irp2* and *iucD*, thus genetically incapable of gaining iron via siderophores salmochelin, aerobactin and yersiniabactin. Besides, *chuA* and *iroN* were absent in all the isolates belonged to groups C and D, respectively. Statistical analysis of the distribution of the genes within the phylogroups showed significant presence of *fecA* in group A, while the presence of *chuA* was related to groups D and E. Other statistically significant relations are presented in details in table 4.

**Table 4.** Distribution of iron acquisition related genes within phylogenetic groups in MPEC and fecal *E. coli* isolates

| Phylogroup | A |  |  | B1 |  |  | B2 |  |  | C |  |  | D |  |  | E |  |  | Unknown |  |  |
| --- | --- | --- | --- | --- | --- | --- | --- | --- | --- | --- | --- | --- | --- | --- | --- | --- | --- | --- | --- | --- | --- |
| Source | M | F | T | M | F | T | M | F | T | M | F | T | M | F | T | M | F | T | M | F | T |
| n | 66 | 16 | 82 | 63 | 35 | 98 | 2 | 0 | 2 | 10 | 1 | 11 | 5 | 5 | 10 | 6 | 3 | 9 | 5 | 0 | 5 |
| *iroN* | *2 | 4 | *6 | 9 | 4 | 13 | - | - | - | *8 | 1 | *9 | - | - | - | - | - | - | *3 | - | *3 |
| *iutA* | *3 | - | *3 | 5 | 1 | 6 | *2 | - | *2 | *9 | 1 | *10 | - | *2 | 2 | - | - | - | - | - | - |
| *fecA* | *65 | *13 | *78 | 57 | *12 | *69 | 1 | - | 1 | 10 | 1 | 11 | 5 | 3 | 8 | *3 | 1 | *4 | 5 | - | 5 |
| *fyuA* | *5 | 3 | 8 | 8 | 3 | 11 | *2 | - | *2 | 3 | - | 3 | *3 | 2 | *5 | - | - | - | *3 | - | *3 |
| *sitA* | 11 | 1 | 12 | *9 | *1 | *10 | 2 | - | *2 | *9 | 1 | *10 | 3 | *3 | *6 | 3 | 1 | 4 | 1 | - | 1 |
| *irp2* | *5 | 2 | *7 | 9 | 4 | 13 | *2 | - | *2 | 3 | - | 3 | *3 | 2 | *5 | - | - | - | *3 | - | *3 |
| *iucD* | *2 | - | *2 | 4 | 1 | *5 | *2 | - | *2 | *9 | 1 | *10 | - | *2 | 2 | - | - | - | - | - | - |
| *chuA* | *- | 1 | *1 | *- | *- | *- | *2 | - | *2 | - | - | - | *5 | *5 | *10 | *6 | *3 | *9 | - | - | - |
| *tonB* | 66 | 16 | 82 | 63 | 34 | 97 | 2 | - | 2 | 10 | 1 | 11 | 5 | 5 | 10 | 6 | 3 | 9 | *4 | - | *4 |

M= MPEC, F= Fecal, T= Total; \* = significant difference ( $p < 0.05$ ); Colored cells = $p < 0.01$ .

### Bacterial survival in raw bovine milk

Eighteen isolates belonging to different genetic patterns (table 5) were cultured in raw milk. According to the results of the bacterial counts up to 8 hours after inoculation, the growth potential of isolates was divided into three groups: no growth, low growth potential (≤ 10^4^ CFU/ml) and high growth potential (> 10^4^ CFU/ml). A significant association (*p*= 0.000) was observed between possession of *fecA* and growth potential in raw milk. Interestingly, all *fecA*+ strains showed a significant growth rate. In contrary, half of the *fecA*-isolates (5/10) could not tolerate the condition of whole milk until the 8^th^ hours after inoculation and were not detected in routine bacterial culture. The remaining *fecA*-isolates showed a lesser growth rate compared with the *fecA*+ members (figure 1). It should be noted that sources of isolates (*p*= 0.165) and IAG scores (*p*= 0.320) had no significant effects on the growth capacity of tested strains.

**Table 5.** Growth potential of *E. coli* isolates of different iron acquisition genotypes in raw bovine milk

| No._Source<br>of isolate | Genotype | Growth<br>potential in raw<br>milk | Viable count at 8h<br>(CFU/ml) |
| --- | --- | --- | --- |
| ***fecA*<sup>+</sup> isolates** |  |  |  |
| 5_M | *fecA_tonB* | High | 10<sup>7</sup> |
| 328_F | *fecA_tonB* | High | 10<sup>7</sup> |
| 47_M | *iroN_fecA_tonB* | High | 10<sup>7</sup> |
| 321_F | *iroN_fecA_fyuA_irp2_tonB* | High | 10<sup>7</sup> |
| 313_F | *iroN_iutA_fecA_sitA_iucD_tonB* | High | 10<sup>6</sup> |
| 19_M | *iroN_iutA_fecA_fyuA_sitA_irp2_iucD_tonB* | High | 10<sup>7</sup> |
| 59_M | *iutA_fecA_fyuA_sitA_irp2_iucD_chuA_tonB* | High | 10<sup>5</sup> |
| 302_F | *iutA_fecA_fyuA_sitA_irp2_iucD_chuA_tonB* | High | 10<sup>7</sup> |
| ***fecA*<sup>-</sup> isolates** |  |  |  |
| 315_F | *tonB* | Low | 10<sup>3</sup> |
| 13_M | *tonB* | No growth | NC at 5h and 8h |
| 6_M | *iroN_tonB* | No growth | NC at 5h and 8h |
| 212_M | *chuA_tonB* | No growth | NC at 8h |
| 420_F | *irp2_tonB* | No growth | NC at 2h and later |
| 409_F | *fyuA_tonB* | Low | $10^4$ |
| 211_M | *sitA_tonB* | Low | $10^2$ |
| 429_F | *sitA_tonB* | Low | $10^3$ |
| 407_F | *chuA_tonB* | Low | $10^3$ |
| 2_M | *iutA_fyuA_sitA_irp2_iucD_chuA_tonB* | No growth | NC at 2h and later |

M= MPEC, F= fecal, NC= not detected in culture; Growth potential was determined based on the viable count at 8h of incubation: 0 CFU/ml= No growth, Low= $10^4 \geq$ , High= $10^4 <$ .

**Figure 1.**
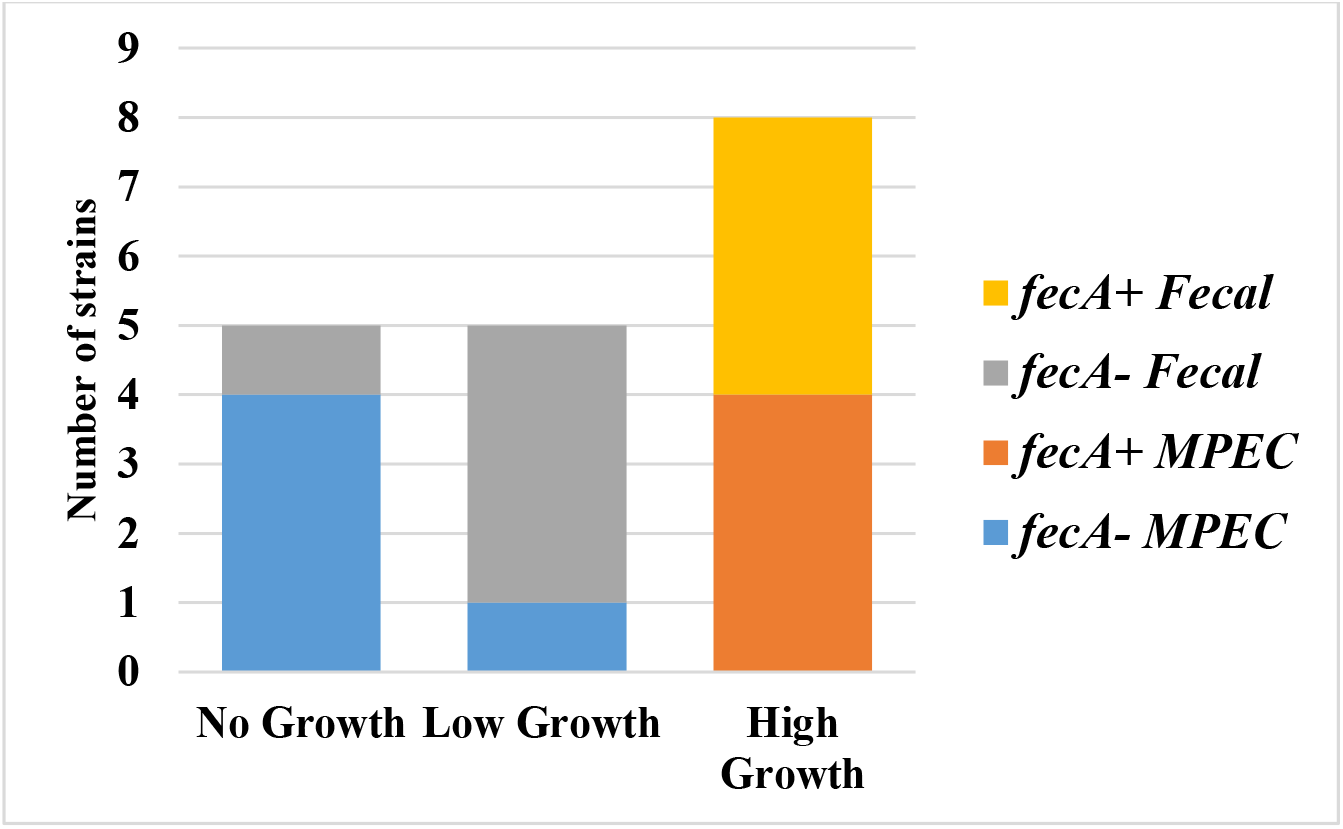
Growth potential of 18 *E. coli* isolates of different iron acquisition genotypes in raw bovine milk.

### Bacterial growth in UHT bovine milk

Twenty-two *E. coli* isolates with different genotypes (table 6) were cultured in UHT bovine milk. Statistical analysis revealed a significant growth potential (*p*= 0.000) for *fecA*+ isolates in UHT milk. This relation was observed at 3, 5 and 8 hours of incubation. Figure 2 and figure 3 graphically represent the results. It is notable that source of strains had no significant effects on bacterial growth (*p*= 0.431).

**Table 6.** Bacterial count of *E. coli* isolates of different iron acquisition genotypes in UHT milk (CFU/ml).

| No._Source of |  | Bacterial counts (CFU/ml) |  |  |  |  |
| --- | --- | --- | --- | --- | --- | --- |
| isolate | Genotype | 0h | 2h | 3h | 5h | 8h |
| *fecA*+ isolates |  |  |  |  |  |  |
| 324_F | *fecA* | 115 | 2185 | 26000 | 4900000 | 945000000 |
| 5_M | *fecA_tonB* | 35 | 1460 | 37500 | 3200000 | 560000000 |
| 68_M | *fecA_tonB* | 95 | 1735 | 32500 | 3500000 | 3287500000 |
| 72_M | *fecA_tonB* | 175 | 2000 | 28375 | 2150000 | 550000000 |
| 328_F | *fecA_tonB* | 130 | 7750 | 448500 | 7700000 | 1105000000 |
| 403_F | *fecA_tonB* | 110 | 650 | 7900 | 610000 | 205500000 |
| 69_M | *iroN_fecA* | 160 | 6985 | 18800 | 9350000 | 2375000000 |
| 47_M | *iroN_fecA_tonB* | 80 | 1875 | 29700 | 4400000 | 1710000000 |
| 310_F | *iroN_fecA_tonB* | 95 | 3850 | 38500 | 3800000 | 835000000 |
| 102_M | *fecA_sitA_tonB* | 280 | 3722 | 32250 | 4250000 | 320000000 |
| 19_M | *iroN_iutA_fecA_fyuA_sitA_irp2_iucD_tonB* | 40 | 670 | 15600 | 1190000 | 448000000 |
| *fecA*- isolates |  |  |  |  |  |  |
| 13_M | *tonB* | 70 | 1810 | 5400 | 23750 | 2740000 |
| 315_F | *tonB* | 10 | 860 | 2790 | 9300 | 94000 |
| 6_M | *iroN_tonB* | 30 | 665 | 3367 | 8950 | 190500 |
| 409_F | *fyuA_tonB* | 205 | 2035 | 8000 | 52000 | 980000 |
| 211_M | *sitA_tonB* | 260 | 8800 | 210550 | 239000 | 30450000 |
| 420_F | *irp2_tonB* | 70 | 2190 | 6900 | 79500 | 6350000 |
| 4201_F | *irp2_tonB* | 50 | 2200 | 6400 | 20100 | 590000 |
| 212_M | *chuA_tonB* | 130 | 3025 | 15800 | 63000 | 2863000 |
| 402_F | *fyuA_irp2_tonB* | 105 | 2320 | 7950 | 58500 | 1100000 |
| 127_M | *iroN_fyuA_sitA_irp2_tonB* | 120 | 1515 | 12300 | 69000 | 14400000 |
| 2_M | *iutA_fyuA_sitA_irp2_iucD_chuA_tonB* | 115 | 700 | 3850 | 15550 | 281000 |
| ***p-value*** |  | 0.606 | 0.847 | **\*0.000** | **\*0.000** | **\*0.000** |

M= MPEC, F= fecal; \* = significant difference ( $p < 0.05$ ).

**Figure 2.**
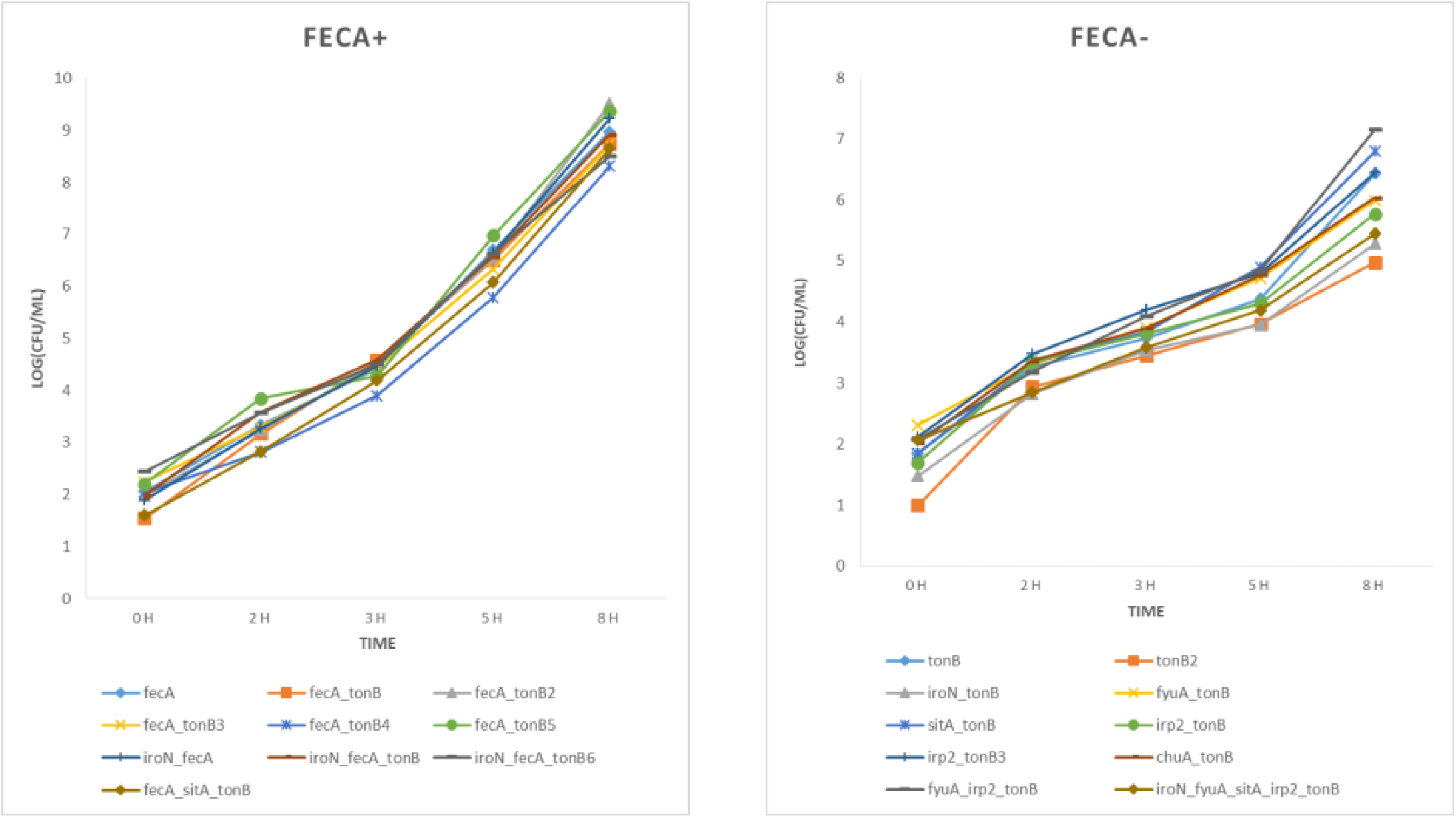
Growth curves of *E. coli* strains in UHT milk by possession of *fecA*.

**Figure 3.**
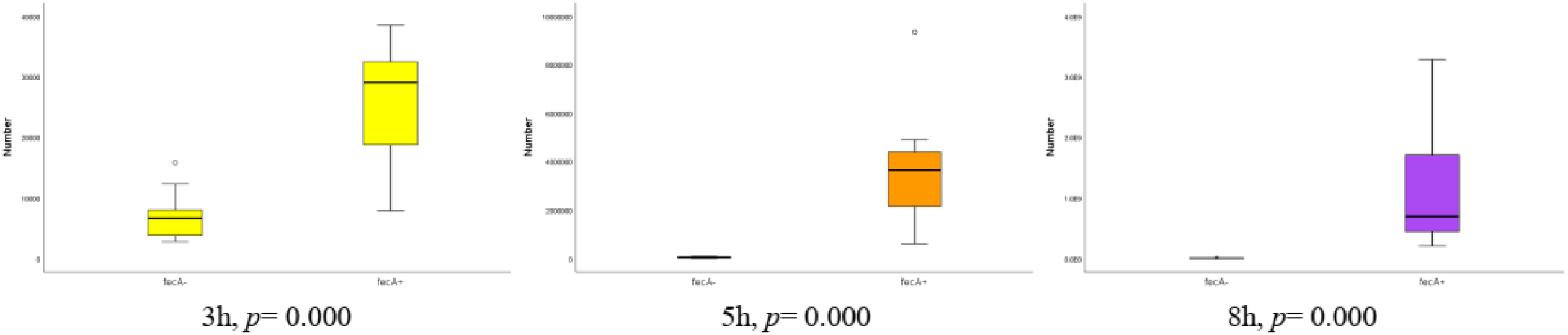
Significant differences in growth rates of *fecA*+ and *fecA- E. coli* strains in 3, 5 and 8 hour of incubation.

### Quantitative analysis of minerals consumption

Significant association was observed (*p* < 0.05) between possession of *fecA* and consumption amount of each of the following minerals: iron (Fe), zinc (Zn), manganese (Mn) and calcium (Ca). Interestingly, *fecA*+ strains consumed less amounts of all minerals compare to *fecA*-isolates. Table 7 and represent results in details.

**Table 7.**
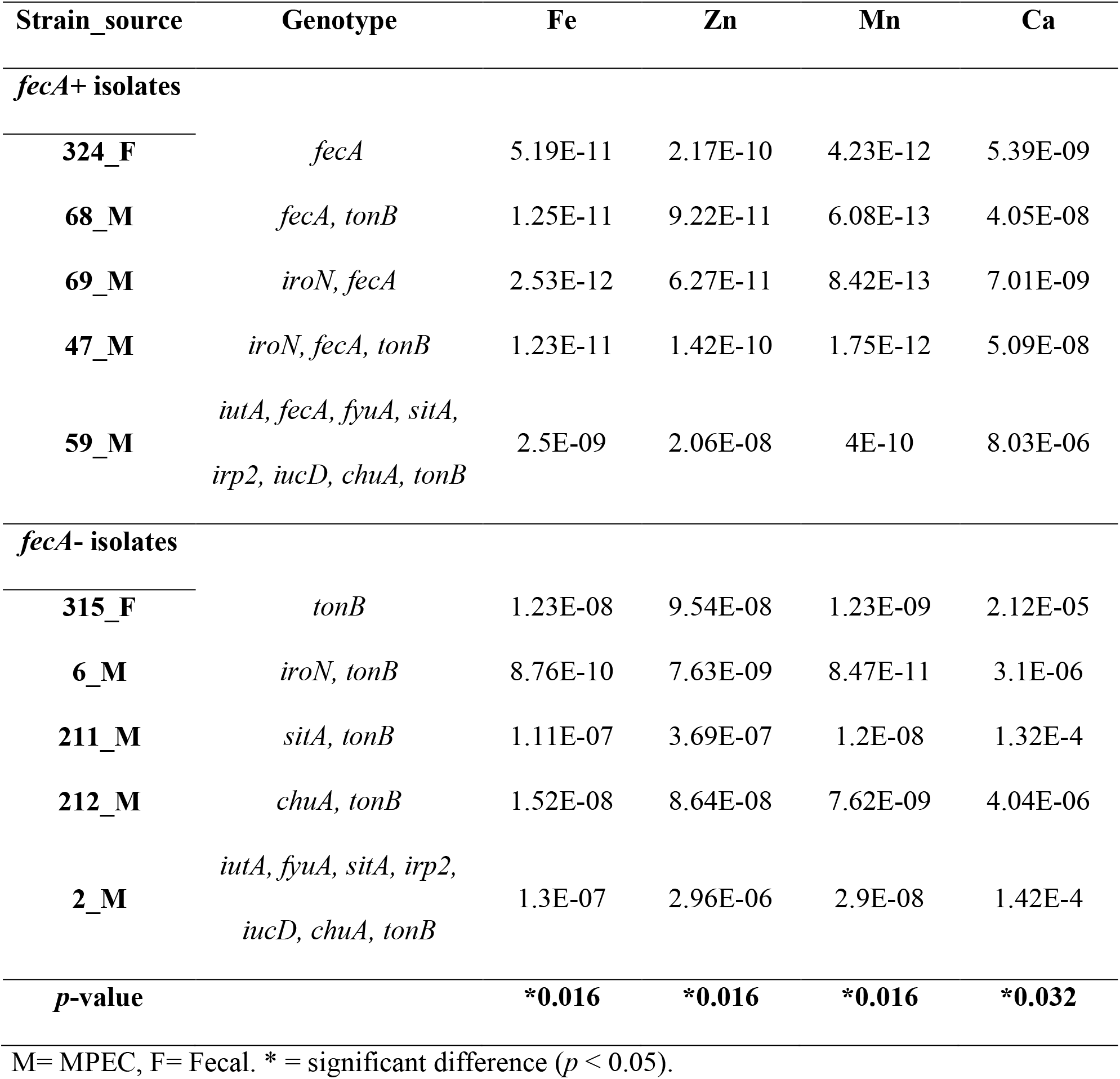
Mineral consumption (ppm/CFU).

| Strain_source | Genotype | Fe | Zn | Mn | Ca |
| --- | --- | --- | --- | --- | --- |
| *fecA*+ isolates |  |  |  |  |  |
| 324_F | *fecA* | 5.19E-11 | 2.17E-10 | 4.23E-12 | 5.39E-09 |
| 68_M | *fecA, tonB* | 1.25E-11 | 9.22E-11 | 6.08E-13 | 4.05E-08 |
| 69_M | *iroN, fecA* | 2.53E-12 | 6.27E-11 | 8.42E-13 | 7.01E-09 |
| 47_M | *iroN, fecA, tonB* | 1.23E-11 | 1.42E-10 | 1.75E-12 | 5.09E-08 |
| 59_M | *iutA, fecA, fyuA, sitA, irp2, iucD, chuA, tonB* | 2.5E-09 | 2.06E-08 | 4E-10 | 8.03E-06 |
| *fecA*- isolates |  |  |  |  |  |
| 315_F | *tonB* | 1.23E-08 | 9.54E-08 | 1.23E-09 | 2.12E-05 |
| 6_M | *iroN, tonB* | 8.76E-10 | 7.63E-09 | 8.47E-11 | 3.1E-06 |
| 211_M | *sitA, tonB* | 1.11E-07 | 3.69E-07 | 1.2E-08 | 1.32E-4 |
| 212_M | *chuA, tonB* | 1.52E-08 | 8.64E-08 | 7.62E-09 | 4.04E-06 |
| 2_M | *iutA, fyuA, sitA, irp2, iucD, chuA, tonB* | 1.3E-07 | 2.96E-06 | 2.9E-08 | 1.42E-4 |
| *p*-value |  | *0.016 | *0.016 | *0.016 | *0.032 |

M= MPEC, F= Fecal. \* = significant difference ( $p < 0.05$ ).

## Discussion

In the present study, we provided some new insights into the usefulness of iron acquisition systems genotyping in pathotype discrimination of MPEC strains. Among various iron acquisition systems investigated in the current study, the significant presence of *fecA* in MPEC was observed and a majority of MPEC belonged to phylogenetic group A. Moreover, we clarified the relations between iron obtaining systems, and growth potential/survivability in raw and UHT bovine milk. Interestingly, the high susceptibility of *fecA-* strains to raw milk environment was observed and the high growth capacity of *fecA*+ strains in both raw and UHT milk was shown in this study. Besides, we investigated the behavior of different *E. coli* strains having different iron acquisition systems to mineral consumption. This study, for the first time provided evidence for a new point of view which is the different nature of *fecA*+ strains in mineral usage, especially iron that is the main subject of the study.

Few recent studies reported a significant presence of ferric citrate outer membrane receptor gene (*fecA*) in MPEC and our findings are in line with them (2–4, 11). Among them, only two studies assessed the effect of *fecA* on phenotypic features. They showed that the presence of *fecA* is required for high growth rate of *E. coli* in milk which is also supported by the current study (3, 4). However, one of the main features which distinguishes our study from the aforementioned ones is that we investigated the phenotypic traits of wild-type strains of *fecA*+ and *fecA-E. coli*. In other words, this is the first study which has evaluated the effect of *fecA* on phenotypic traits of *E. coli* on larger scale (numbers of tested strains) and using clinical isolates.

In contrast to limited surveys on *fec* locus in MPEC, several reports have been recorded about other iron acquisition systems with focusing on famous siderophores. In fact, the predominant approach of these studies were to investigate the presence of well-known virulence genes of diarrheagenic *E. coli* (DEC) and ExPEC pathotypes such as genes encoding toxins, pili, fimbriae, metal scavenging systems and etc. However, the most hinted iron acquisition genes in previous studies are *iucD*, *iroN*, *fyuA* and *irp2*. Moreover, the data about *chuA* can be extracted from the results of phylogenetic grouping conducted based on the Clermont’s procedures. Prevalence of the genes *iucD, irp2* and *chuA* are recorded in ranges of 0-16.7% (12, 13), 2.5-26.4% (13, 14) and 5.7-35.6% (15, 16), respectively by other researchers, and our findings (*iucD*= 10.8%, *irp2*= 15.9% and *chuA*= 8.3%) are situated in the ranges. Nevertheless, presence of *fyuA* (15.3%) and *iroN* (14%) among our isolates were higher than previous records (2.5% and 3.5%, respectively) (12, 14). Unfortunately, our results of iron acquisition genotypes are incomparable with most of the previous studies, as no focus on iron uptake systems can be observed among them.

In contrary to the general belief that ExPEC strains are rich in iron acquisition systems involved in pathogenesis, we showed that most bovine MPEC and fecal strains are lacking complex iron uptake genotypes and most only harbored *fecA_tonB* (MPEC) and *tonB* (fecal strains). This, could be a distinguishing factor for MPEC from other ExPEC like Uropathogenic *E. coli* (UPEC) and Avian pathogenic *E. coli* (APEC) which mostly benefit from the simultaneous presence of different iron acquisition systems including siderophores (17, 18). A possible explanation for such tendency in MPEC could be the fact that the siderophores are energetically expensive and could be hijacked by other bacteria in the environment (4, 19). In explaining the high frequency of *fecA-tonB* in MPEC isolates, we can also pay attention to the role of citrate as an alternative energy source. *E. coli* possesses all of the enzymes necessary for citrate metabolism, so citrate can be utilized, even if the imported amount is small (20).

*E. coli* populations are not randomly distributed in different phylogroups and members of a same phylogenetic group are more likely to have a common genotypic and phenotypic features, same origin and ability to induce diseases (21). The current study and previous works report MPEC isolates belong mostly to groups A and B1 which suggests derivation of MPEC from commensal ancestors (11, 22, 23). Unlike most APEC and UPEC isolates which are members of B2 and D phylogenetic groups, this study and other investigations indicates that the most of MPEC do not share same lineages similar to other ExPEC (17, 24). Although the associations between iron acquisition genotype and phylogenetic groups are not clearly elucidated, there are studies that have mentioned some possible relations. For instance, there were significant relations between *iutA* and group B2 (25), *iucD* and groups B2 and D (26), and *irp2* and group A (15) that our results also support these findings. Furthermore, the significant associations of *fecA* with group A and *chuA* with groups D and E were detected in the current study.

Most previous studies on MPEC have relied on the assessments of virulence features, host-pathogen interactions, and fitness elements for proliferation in milk and/or udder (4, 15, 22, 23, 27, 28). The latter, has been evaluated from various aspects such as effects of genes involved in nucleotide and amino acid biosynthesis, *fecIRABCDE* locus and alteration in LPS on growth potential of *E. coli* in milk (3, 4, 29). Among different recorded results, the vital role of *fec* locus in high growth rates of *E. coli* in milk is in agreement with our findings (3, 4). However, the other distinguishing feature of the current study is that the growth behavior of *E. coli* in milk was evaluated from a new perspective of owning different iron acquisition systems. It was observed that only *fecA* and no other iron acquisition systems, played a significant role in both *E. coli* survival and growth potential in milk and sources of the isolates had no considerable effect on these attributes. All the *fecA*+ isolates were fast-growing while half of the *fecA-* strains could not even tolerate the raw milk milieu. It could be due to the more susceptibility of *fecA-* isolates to be cleared by immune cells of udder and milk (3). Since we observed *fecA-* isolates could not effectively survive and grow in raw milk, mastitis development by *fecA*-strains needs to be clarified in future investigations. As noted before, the significant differences in growth rates of *E. coli* strains had begun from the 3 hour after inoculation in UHT milk. Therefore, a lesser duration of incubation in future studies assessing the growth potential of MPEC is proposed. It seems that any period longer than three-hour of incubation is sufficient to examine the growth capacity of *E. coli* in milk.

The relations among genotypes, growth potential of the strains in milk and consumption amount of iron were also clarified. Interestingly, it was found out the fast-growing strains which all had *fecA*, significantly consumed less amounts of iron per colony-forming unit (CFU) in comparison with *fecA-* strains even the ones with the ability of gaining iron from other systems. We believe that attention to other aspects of bacterial metabolism like optimal energy consumption can be a clue for clear explanations why two groups of *E. coli* strains (*fecA*+ and *fecA*-) have different growth rates in milk and consume different amounts of iron. Based on the Fenton reaction which is capable of generating both hydroxyl radicals and higher oxidation states of the iron, the more iron bacteria gain, the more radicals they should face (30). Thus, *fecA*+ strains which acquire less amounts of iron may have the benefit of saving energy from less neutralization of radicals while accessing to citrate as an alternative energy source. Along with genes only involved in iron acquisition, we also screened isolates for possession of *sitA* which is responsible for both iron and manganese transportation (31). From this point of view, we also measured consumption amounts of manganese. Unexpectedly, results were as the same as the relation between iron consumption and *fecA*. Since the data of consumption amounts of zinc and calcium by cultured strains were available, a comparison was also taken to identify any relations among genotypes of the strains and minerals consumption. Results were similar to the relations between iron and manganese consumption and *fecA* which revealed a new behavior of *fecA*+ strains in economic mineral acquisition.

## Conclusion

According to the results, it seems that different *E. coli* strains do not share similar necessities. Specifically, genetic variations in terms of possessing iron acquisition systems were observed in MPEC, despite the most of MPEC strains shared common phylogenetic lineages A or B1. Analysis of the genetic associations with phenotypes revealed that among different iron acquisition systems iron uptake through ferric citrate plays a significant role in the behavior of *E. coli* in milk. The current study lit up a new aspect of the nature of *E. coli* and its consequent behaviors. However, we believed that more investigations of mentioned points by attention to other bacterial traits such as optimal energy consumption and metabolomics are necessary for a better comprehension of *E. coli* pathogenicity.

## Material and Methods

### Sampling and bacterial isolates

Clinical mastitis milk samples were collected aseptically from four industrial dairy farms during April 2017 to July 2019. Milk samples were cultured and interpreted according to recommendations of National Mastitis Council (32). To isolate *E. coli* causing mastitis strains, only pure cultures of lactose positive bacteria with relatively high load of similar colonies were chosen for further identification. Conventional biochemical tests were carried out in order to confirm bacterial species (33).

To compare the pathogenic strains with commensals, some samples were also taken from feces. Fecal samples were taken from cows suffering from clinical mastitis. Samples were cultured on MacConkey agar plates. After 24 hours incubation at 37°C, any lactose positive colonies were investigated in order to isolate *E. coli* strains. Confirmation was based on the results of conventional biochemical tests (33).

A total of 217 confirmed *E. coli* isolates (mastitis isolates= 157, fecal isolates= 60) were cryopreserved for further investigations.

### DNA extraction

Boiling procedure was applied for DNA extraction (34). Briefly, a loop of cultured bacteria was mixed with 500 μl sterile distilled water and was suspended by vortexing for 30 s. Then, the suspension was boiled in 98°C for 10 min and was placed on ice for another 10 min. Finally, it was centrifuged in 8000 ×g for 5 min. The supernatant was collected and used as template DNA.

### Molecular profiling of iron acquisition systems

Presence of nine different genes related to diverse iron acquisition systems were investigated through the recently developed multiplex PCR assays which were conducted in three panels. Each panel is capable of simultaneously screening three genes and they are as follows: panel 1 *iroN* (salmochelin siderophore receptor), *iutA* (aerobactin siderophore receptor) and *fecA* (ferric citrate receptor); panel 2: *fyuA* (yersiniabactin siderophore receptor), *sitA* (ferrous iron/ manganese transporter substrate-binding) and *irp2* (biosynthesis of siderophores yersiniabactin); and panel 3: *iucD* (biosynthesis of the siderophores aerobactin), *chuA* (haem receptor) and *tonB* (energy transducer) (34).

### Determination of phylogenetic groups

Phylogenetic groups were determined based on the method described by Clermont et al., (21). The method is capable of assigning the *E. coli* isolates to groups: A, B1, B2, C, D, E, F and clades II to V. However, there was a small portion of isolates categorized as unknown.

### Phenotypic assays

All the phenotypic assays (bacterial survival in raw bovine milk, bacterial growth in commercial UHT bovine milk, and quantitative determination of mineral consumption) were culture-based which had similar procedures in steps of preparation of inoculum and culture.

### Inoculum preparation

*E. coli* strains were first cultured on MacConkey agar to obtain single colony. After overnight incubation at 37°C, a single colony was inoculated in 3 ml of BHI broth. Then, the culture was incubated at 37°C until its turbidity was adjusted to a 0.5 McFarland standard (OD= 0.08-0.12 at 625 nm). Thereupon, bacterial cultures were diluted and suspensions at a concentration of 10^4^ CFU/ml were achieved.

### Bacterial survival in raw bovine milk

A total of 18 *E. coli* isolates were selected based on their genotypes (table 5) to evaluate any associations between genotypes and the ability of strains to survive in raw bovine milk. One of the fundamental and very first step for mastitis development by *E. coli* is prevailing over harsh barriers exist in milk and udder (phagocytic cells, antibodies, complements, etc.) to survive. Therefore, raw milk was considered as culture media to imitate natural conditions.

Raw milk from healthy dairy cows were collected aseptically one day prior to inoculation. To examine the microbial load, 500 μl of each samples were cultured on blood agar and incubated at 37°C. Next, five milk samples with no microbial growth on blood agar cultures were chosen and mixed together for inoculation of prepared cultures. According to the procedure described above, 25 μl of the final inoculum (10^4^ CFU/ml) was prepared and added to 5 ml of bovine raw milk. Bacterial survival was determined by standard plate count method at 0, 2, 3, 5 and 8 hours after inoculation. For this purpose, enumeration of colonies by culturing of 100 μl of each samples on TSA plates was applied. The procedure was conducted in double.

### Bacterial growth in commercial UHT bovine milk

Another vital step for *E. coli* to develop mastitis is propagation in relatively high number. In order to assess effect of genotype on the growth rate of *E. coli* in a poor-iron environment, ultra-high temperature milk (UHT) was chosen as culture media. UHT milk has also the advantage of being similar to natural media in biological contents including protein and fat. Moreover, the inhibitory effects of complement and phagocytes are neutralized in UHT milk which increases the precision of evaluating the relations among iron, genotypes and growth rates. To analyze bacterial growth rates in UHT milk, the procedure bellow was followed:

A total of 22 *E. coli* isolates were selected based on their genotypes (table 6) and commercial UHT milk was obtained. A volume of 5 ml of milk was inoculated with 25 μl of bacterial suspension (10^4^ CFU/ml) as described before. The inoculated milk was dispensed in five 1.5 ml microtubes (1 ml per tube). Microtubes were incubated at 37°C. Bacterial counts were determined by serial dilutions and duplicate plating on TSA plates at 0, 2, 3, 5 and 8 hours after inoculation.

### Quantitative determination of mineral consumption

To determine any probable relations among mineral consumption and genotypes, 10 *E. coli* isolates were chosen based on the genetic patterns related to iron acquisition systems (table 7). According to the procedure described in the previous section, isolates were cultured in UHT milk and bacterial counts were determined at the time of the inoculation and 8 hours after incubation. After 8 hours, samples were centrifuged in 4000 ×g for 6 min to separate bacterial cells. The supernatants were collected for quantitative analysis of iron (Fe), zinc (Zn), manganese (Mn), and calcium (Ca) contents by inductivity coupled plasma-optical emission spectroscopy (ICP-OES) (Spectro Arcos, Germany). A part of the UHT milk which was used as culture media, kept intact and its mineral contents were also measured by ICP-OES. The consumption amounts of minerals per CFU for each isolate were calculated using the following formula:

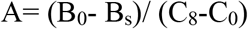

Where:

A: Mineral consumption per CFU for sample (ppm/CFU)
B_0_: Concentration of the intended mineral in the control milk sample (ppm)
B_s_: Concentration of the intended mineral in the supernatant of the cultured milk sample (ppm)
C_8_: Viable bacterial count at 8 hours of incubation (CFU)
C_0_: Viable bacterial count at the time of milk inoculation (CFU)

### Statistical analysis

The data was statistically analyzed using SPSS version 16.0 (SPSS Inc., Chicago, USA). The chi-square test was applied to compare categorical variables. Mann-Whitney also used for analyses related to bacterial growth rate in UHT milk and mineral consumptions. *P*-value < 0.05 was considered to be statistically significant.

## Acknowledgments

This work is supported by Ferdowsi University of Mashhad under the project number 48474. The authors would like to thank Sara Arab (Shahid Beheshti University, Tehran) for kind helps in statistical analysis.

